# ACME dissociation: a versatile cell fixation-dissociation method for single-cell transcriptomics

**DOI:** 10.1101/2020.05.26.117234

**Authors:** Helena García-Castro, Nathan J Kenny, Patricia Álvarez-Campos, Vincent Mason, Anna Schönauer, Victoria A. Sleight, Jakke Neiro, Aziz Aboobaker, Jon Permanyer, Marta Iglesias, Manuel Irimia, Arnau Sebé-Pedrós, Jordi Solana

## Abstract

Single-cell sequencing technologies are revolutionizing biology, but are limited by the need to dissociate fresh samples that can only be fixed at later stages. We present ACME (<u>AC</u>etic-<u>ME</u>thanol) dissociation, a cell dissociation approach that fixes cells as they are being dissociated. ACME-dissociated cells have high RNA integrity, can be cryopreserved multiple times, can be sorted by Fluorescence-Activated Cell Sorting (FACS) and are permeable, enabling combinatorial single-cell transcriptomic approaches. As a proof of principle, we have performed SPLiT-seq with ACME cells to obtain around ∼34K single cell transcriptomes from two planarian species and identified all previously described cell types in similar proportions. ACME is based on affordable reagents, can be done in most laboratories and even in the field, and thus will accelerate our knowledge of cell types across the tree of life.

## Introduction

Biology is undergoing a paradigm shift due to the introduction of single-cell sequencing methods [1-5]. The cell is the fundamental unit of biological systems and studying thousands of them individually allows reconstruction of the cellular diversity and dynamics formerly blended into bulk tissue samples. For instance, single-cell transcriptomics (or scRNA-seq) allows measurement of the expression of thousands of mRNAs from potentially hundreds of thousands of individual cells. The mRNAs of each cell are indicative of the cell type or state, and allow biological questions to be addressed at a new level of integration and detail. From the sequencing of a single cell in 2009 [6] we have seen year on year exponential increases in the number of cells that can be sampled by scRNA-seq [7]. Using these methods, scientists have already profiled a broad taxonomic range of different animals, classified their cell types, profiled their gene expression patterns, and begun to reconstruct their cell differentiation lineages. Single cell transcriptomics has been already used in very diverse animal groups, including sponges [8, 9], cnidarians [10, 11], planarians [12-16], nematodes [17, 18], arthropods [19-22], ascidians [23] as well as extensively in vertebrates [24-31].

Currently, the most popular methods are based on nanodroplet barcoding: encapsulating single cells with oligonucleotide barcodes in nanolitre droplets [32, 33]. One of the most promising recent developments involves employing combinatorial barcoding techniques [17, 34], which use the cells themselves as reaction chambers. These approaches barcode cellular mRNAs through successive rounds of mixing and pooling the cell population such that the probability of two cells receiving the same barcode combination is minimised. The implementation of combinatorial barcoding methods allows the generation of datasets containing millions of cells from different samples [25].

One major technical hurdle of single-cell transcriptomic approaches is the lack of a cell dissociation method that simultaneously fixes the cells and preserves mRNAs. Typically, dissociation is done in live cells and relies on enzymatic (*e*.*g*. trypsin, papain or similar) or mechanical (e.g. dounce homogenisation) approaches [35], which introduce dissociation artefacts and cellular stress on the samples [36-38]. Dissociated cells are stripped from their extracellular context and washed, incubated, centrifuged, stained and often sorted by FACS while still alive, which changes their gene expression patterns [36-38]. Preservation can only take place hours after the beginning of the experiment, but this time suffices for the activation of stress responses [36]. One alternative approach is obtaining single-cell transcriptomic data from nuclei [39], as these can be extracted from frozen tissue [25]. However, this approach eliminates the majority of mature mRNAs, as these concentrate outside the nucleus. The introduction of a method that simultaneously fixes and dissociates cells, preserving their RNA, is a critical need of the single cell transcriptomic field.

To overcome the limitations of live cell dissociations, we have developed ACME dissociation. Our protocol is based on a 19^th^ century dissociation protocol – often called “maceration” – with modifications to make it compatible with modern single-cell transcriptomics. The maceration procedure was first used by Schneider in 1890 [40]. It was then used throughout the 20^th^ century to dissociate cells of animals such as cnidarians [41] and planarians [42] and observe them under the microscope, but has now largely lapsed into disuse.

In its original form, the maceration solution simply consisted of acetic acid and glycerol dissolved in water. Baguñà and Romero added methanol, as it speeds up dissociation [42]. Our protocol uses acetic acid and methanol, together with glycerol, dissolved in water. This solution produces fixed single cells in suspension with high integrity RNAs. Conveniently, we show that ACME-dissociated cells can be cryopreserved using DMSO [43] at different points throughout the process, with little detriment to their recovery and RNA integrity. As a proof of principle, we combined ACME to an optimised version of SPLiT-seq [34], including 4 rounds of barcoding, and were able to profile 33,827 cells from two different planarian species, *Schmidtea mediterranea* and *Dugesia japonica*, in a single run. We recover all *S. mediterranea* cell types from a previous study [13], at comparable proportions, showing that ACME dissociation does not introduce biases in cell type composition. Furthermore, we describe for the first time the single-cell transcriptome of *D. japonica*, opening the study of cell type evolution in this clade. Thus, in combination with SPLiT-seq or other approaches, ACME dissociation is a robust method to obtain high-quality single-cell transcriptomic data from fixed cells.

## Results

### ACME dissociation produces fixed cells with preserved morphology that can be visualised by cell cytometry

ACME dissociation takes place in ∼1h (Figure 1A). We immerse ∼10-15 adult *S. mediterranea* individuals or a similar amount of other tissue (representing ∼100 µl of biological material) in 10 mL of ACME solution. An optional washing step in N-Acetyl-L-Cysteine (NAC) prior to ACME dissociation helps remove mucus [44] (See Supplementary Note 1). Once animals are in ACME solution, they are shaken for 1h at room temperature, with occasional pipetting up and down of the solution to aid dissociation. We then collect the cells by centrifugation to remove the maceration solution (Figure 1A), and wash the pellet in a PBS solution containing 1% BSA at 4°C. We perform a second centrifugation as a subsequent cleaning step and, finally, resuspend the cells in the same buffer solution (Figure 1A). After this step, cells must be kept in PBS/1% BSA solution in cold conditions (i.e. on ice).

**Figure 1:**
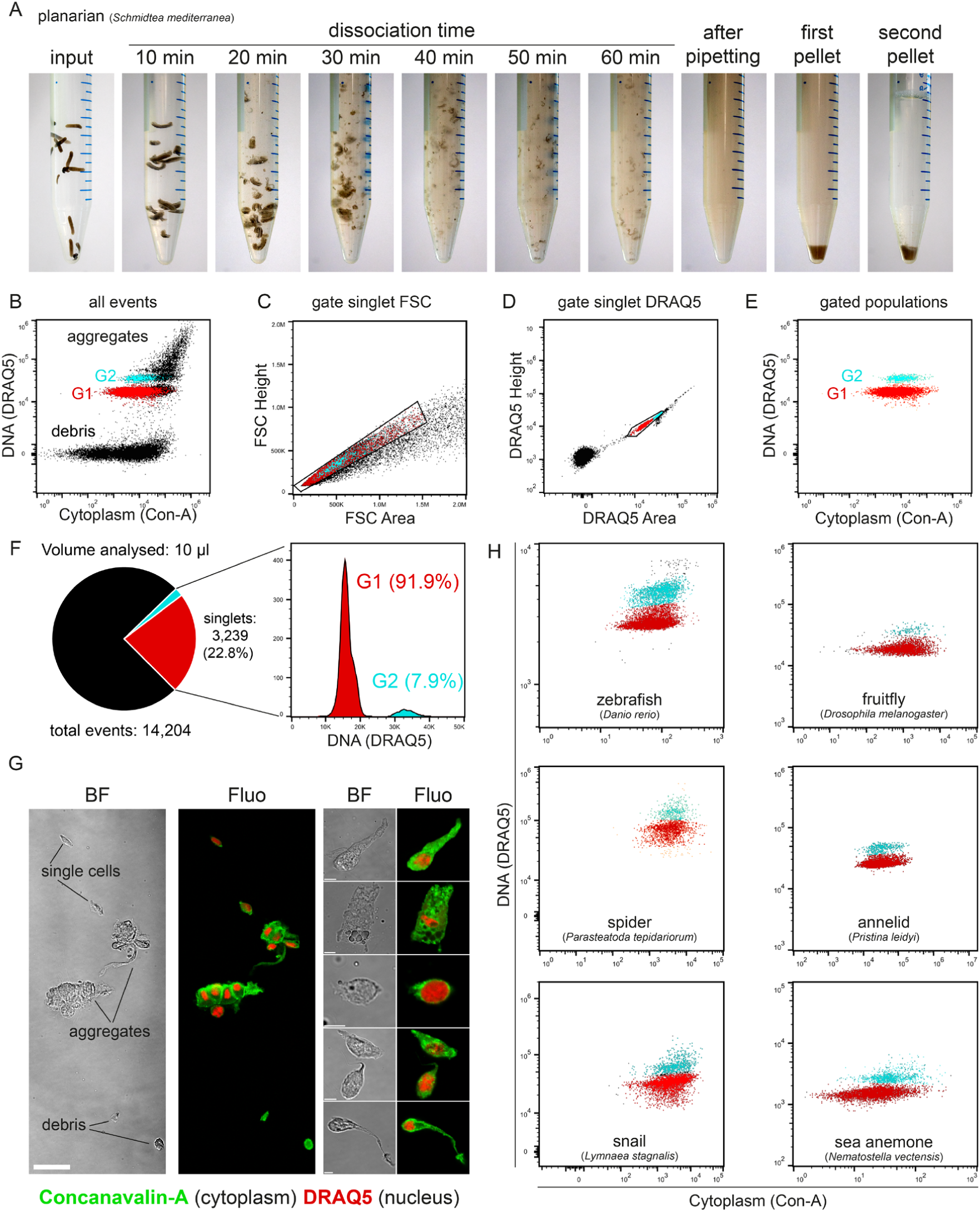
ACME dissociation, a species-versatile cell dissociation method. **A:** Whole dissociation process for the planarian *Schmidtea mediterranea*. From left to right: live worms used as input in water, ACME dissociation reaction after 10-60 minutes, cell suspension after final pipetting, pellet after first centrifugation, pellet after second centrifugation in PBS 1% BSA. **B-E:** Flow cytometry profiles of *S. mediterranea* ACME-dissociated cells stained with DRAQ5 (nucleus) and Concanavalin-A (cytoplasm) ungated (B) and after gating singlets by FSC (C) and DRAQ5 (D) Area vs. Height, resulting in clean G1 (red) and G2 (cyan) populations **F:** Relative proportion of singlets in a typical *S. mediterranea* ACME cell dissociation, corresponding to 22.8% of the total events, and histogram of their DNA content (linear scale), showing the relative proportions of G1 and G2 cells. **G:** Bright field (BF) and confocal fluorescence (Fluo) microscopy images of *S. mediterranea* ACME-dissociated cells stained with Concanavalin-A and DRAQ5, showing single cells, aggregates and debris (left) and details of different cell types with well-preserved morphology (right). Scale bars are 50 µm (left) and 5 µm (right). **H:** Flow cytometry gated profiles of ACME-dissociated cells from different organisms: zebrafish 1-day embryos (*Danio rerio*), fruitfly 3^rd^ instar stage larvae (*Drosophila melanogaster*), spider stage 7 embryos (*Parasteatoda tepidariorum*), annelid adults (*Pristina leidyi*), snail larvae (*Lymnaea stagnalis*), and sea anemone juveniles (*Nematostella vectensis*).

In order to visualise cells by flow cytometry, we stain fixed cells with DRAQ5 (nuclei) and Concanavalin-A conjugated to Alexa Fluor 488 (cytoplasm). DRAQ5 is a far-red emitting, anthraquinone compound that stains DNA. Concanavalin-A is a lectin that binds carbohydrates present in internal cell membranes. Since ACME cells are permeabilised, we find Concanavalin-A staining throughout the cytoplasm. These staining conditions allow us to visualise the cells by flow cytometry, revealing several cell populations (Figure 1B). Both G1 and G2 populations are readily visible, corresponding to cells in different phases of the cell cycle. G2 corresponds to what planarian FACS protocols typically refer to as the ‘X1’ population [45]. Like other dissociation protocols, ACME also produces a large quantity of cellular debris, with cytoplasm staining but without DNA (Figure 1B). Undissociated cell aggregates are also visible, with higher levels of DNA and cytoplasm staining (Figure 1B). Performing a singlet filter by cell cytometry, using FSC (Figure 1C) and DRAQ5 (Figure 1D) area vs height completely removes the aggregates and debris. We also gate Concanavalin-A and DRAQ5 positive cells to obtain clear G1 and G2 populations (Figure 1E).

Typically, we resuspend one dissociation reaction in 1mL of buffer. Analysis of 10 μL of such reactions reveals thousands of singlet cells (Figure 1F). Thus, a reaction with 10 adult *S. mediterranea* generates well over 100K singlet cells that can be FACS sorted. The relative proportions of G1 vs G2 cells are similar to those described in planarians by enzymatic approaches [45] (Figure 1F, right). ACME dissociated cells exhibit well-preserved morphology under microscopic observation (Figure 1G), as this was the original purpose of the maceration technique [40-42].

We successfully dissociated several animals, including zebrafish embryos, fruitfly larvae, spider embryos, annelid adults, snail larvae, and juvenile sea anemones using the same protocol with minimal alterations (Figure 1H). This includes organisms belonging to diverse major metazoan lineages and with a broad range of terrestrial, freshwater and marine habitats. Therefore, ACME dissociation is a versatile method that can be used in markedly different animal models. However, ACME solution dissociates soft tissues and cannot penetrate hard shells, chorions or vitelline membranes. We dechorionated zebrafish embryos using standard protocols and ruptured the cocoons and vitelline membranes that encapsulate spider and snail embryos. This can be done after embryos are placed in the ACME solution, using forceps under the scope or with short pulses of homogenisation. Soft-bodied animals like planarians completely dissociate with minimal mechanical forces (shaking and pipetting up and down) but other animals such zebrafish or cnidarians benefit from stronger mechanical dissociation using a combination of homogenisation and dissection.

### ACME dissociated cells can be cryopreserved multiple times and retain high integrity RNAs

Currently, single cell dissociation protocols typically rely on enzymatic (e.g. trypsin) dissociation. One disadvantage of enzymatic approaches is that cells can only be cryopreserved after dissociation, typically by FACS sorting them into methanol-[46] or DMSO-[43] containing solutions. However, cells have already been out of their cellular context for several hours. Apart from biological effects, this also imposes logistical restrictions: cell dissociation needs to be done in close proximity to a single-cell transcriptomic facility or at least, a FACS facility. This logistical restraint renders single-cell analysis of specimens collected in remote sampling areas, or that are difficult to culture in the laboratory, extremely challenging.

Our protocol comprehensively solves this problem, as ACME fixes cells immediately. Furthermore, we found that ACME-dissociated cells can be easily cryopreserved at several steps of the process by freezing them in a PBS solution containing BSA (1%) and DMSO (10%) [43]. To test this, we compared ACME-dissociated cell populations after several freezing steps (Figure 2A). We analysed by flow cytometry cell populations right after dissociation (Figure 2B), and after freezing and thawing the cells (Figure 2C). We also FACS sorted ACME-dissociated cells resulting in an 85-90% enrichment of G1 and G2 cells. We compared FACS sorted cells directly after sorting (Figure 2D) with FACS sorted cells cryopreserved again after sorting (Figure 2E). This shows that ACME-dissociated cells can be subjected to several rounds of cryopreservation without altering their cytometry profiles. To test the resistance to freezing/thawing cycles, we subjected ACME-dissociated cells to 5 freeze/thaw cycles and analysed the resulting populations (Figure 2F). We found no differences in the cytometry profiles of ACME-dissociated cells after multiple steps of cryopreservation, showing that ACME is a robust and convenient method to obtain and preserve dissociated cells.

**Figure 2:**
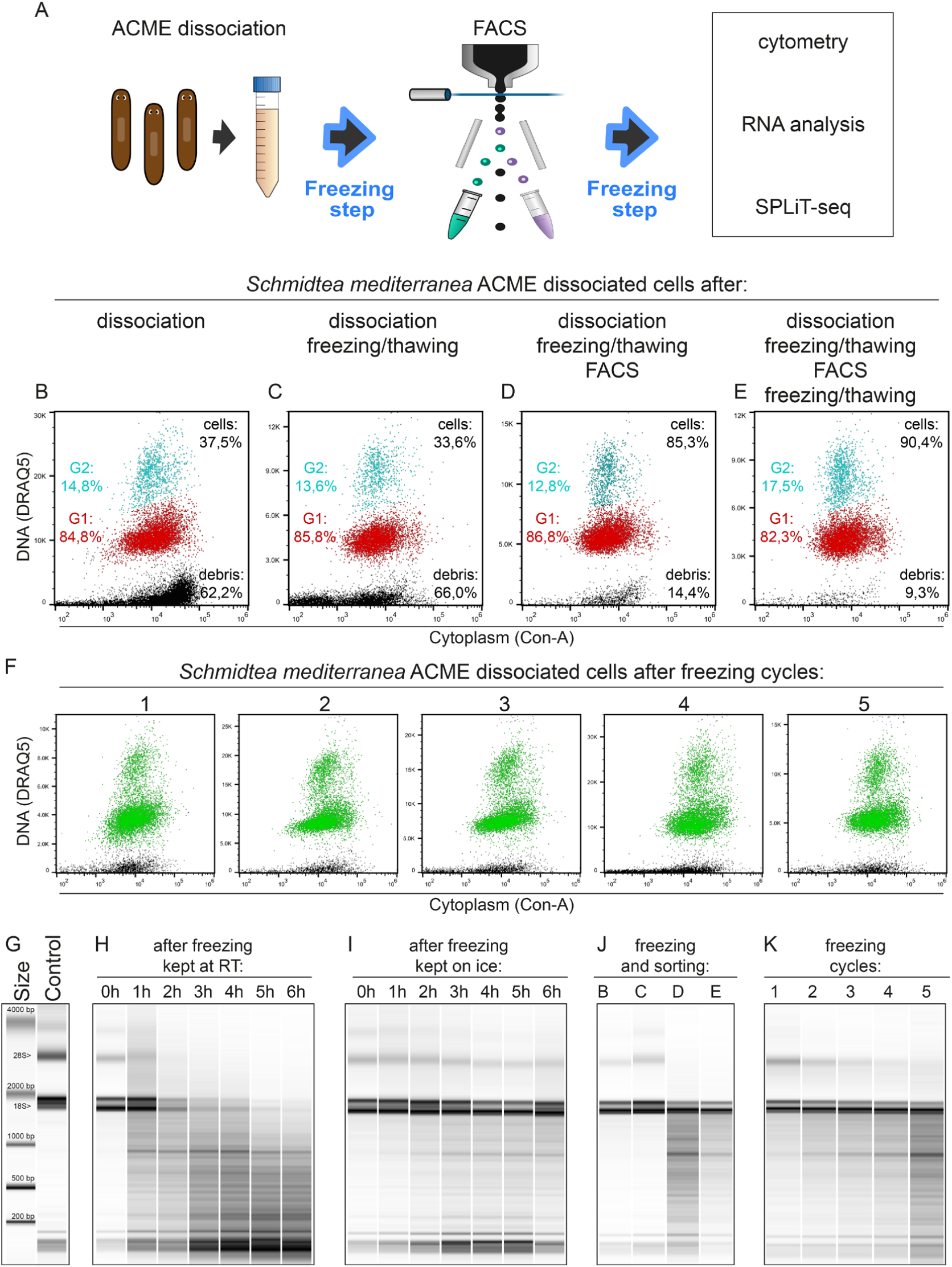
Cryopreservation and RNA integrity of ACME-dissociated cells. **A**: Experimental workflow of ACME dissociation for single-cell RNA transcriptomics. **B-E:** Flow cytometry profiles of singlet (FSC) *Schmidtea mediterranea* ACME-dissociated cells, stained with DRAQ5 (DNA) and concanavalin-A (cytoplasm), directly after dissociation (B), after a first freezing step (C), after FACS (D) and after a second freezing step (E). DRAQ5 scales are shown in linear values and differ due to the freezing steps. Aggregates are gated out by FSC. Percentages relative to the number of total singlets are shown in black for debris and cells. G1 (red) and G2 (blue) percentages refer to these population proportions. G1 and G2 proportions do not greatly vary, but FACS sorting effectively enriches these populations. **F**: Flow cytometry profiles of singlet (FSC) *S. mediterranea* ACME-dissociated cells after 1 to 5 freezing cycles. DRAQ5 and concanavalin A positive cells are shown in green and debris in black. Scale and gating conditions as in B-C. **G-K:** Bioanalyzer profiles of RNA samples. As a control sample, we used RNA from worms dissociated directly in Trizol (G). A size ladder is displayed and the two major RNAs (18S and 28S) are indicated. Time and temperature-dependent RNA degradation were tested keeping samples at room temperature (H) or on ice (I) for 6 hours. RNA integrity along the protocol was tested for the conditions described in B-E (J), showing partial degradation after FACS. RNA integrity was tested after 1 to 5 freeze/thaw cycles cycles (I).

We next tested if RNAs are well preserved in ACME-dissociated cells, a critical requirement for single-cell transcriptomics. We added NAC to the ACME solution as it provides reducing conditions that protect RNA from degradation [44] (See Supplementary Note 1). We found that the major factors affecting RNA quality in ACME-dissociated cells are time and temperature. To test this, we extracted RNA from ACME-dissociated cells, including one freezing step, and compared them to RNAs from undissociated planarians (Figure 2G). To test RNA degradation at room temperature, we incubated ACME-dissociated cells in PBS containing 1% BSA for several hours (Figure 2H). We observed that RNA integrity after ACME dissociation progressively declines over time at room temperature. Keeping cells on ice effectively prevents this effect (Figure 2I). Therefore, cold conditions are sufficient to safeguard mRNA for ongoing work. FACS sorted cells also show some RNA degradation (Figure 2J), as the total time required for our staining and FACS sorting is about ∼3-6 hours. Repeated freeze/thaw cycles also negatively impact RNA integrity (Figure 2K), but do not result in complete degradation. These results show that ACME-dissociated cells can be cryopreserved multiple times with little detriment to recovery and RNA integrity. Therefore, ACME dissociation is a convenient method to obtain and cryopreserve large sample sets and to obtain single cell suspensions of animals difficult to culture in the lab, or directly from the wild.

### Single-cell transcriptomic analysis of ACME-dissociated cells recovers expected cell type composition

To test the potential of ACME-dissociated cells in combinatorial barcoding protocols we performed a species mixing experiment (Figure 3A) using SPLiT-seq [34] to profile two different planarian species: *S. mediterranea* and *Dugesia japonica*. Briefly, pools of cells are split equally into wells, labelled with different reactions, and then pooled again (Figure 3A, Supplementary Figure 1A). After four barcoding rounds the probability of any two cells receiving the same barcode combination is minimised. The SPLIT-seq protocol consists of an in-cell retrotranscription (RT) and two rounds of oligo ligations, with the fourth barcode introduced in the sub-library amplification step. We modified the original SPLiT-seq protocol to make it compatible with ACME-dissociated cells. First, we eliminated the formaldehyde fixation included in the original protocol as ACME-dissociated cells are already fixed by the acetic acid. Furthermore, we eliminated random hexamer RT oligos, as these could result in excessive rRNA recovery, as these are highly abundant in the cytoplasm of all cells. In our configuration, we used 48 RT poly-dT barcodes and 96 ligation barcodes in each of 2 rounds of ligation, and 3 sub-libraries (Figure 3A), which together generate 1.3 million possible barcode combinations. The barcodes are concatenated in the terminal part of the resulting cDNA sub-library, whereas the other end contains the mRNA sequence (Figure 3A, Supplementary Figure 1B). To minimise collisions (cells receiving the same combination of barcodes) it is recommended to use less than 5% of the number of possible combinations. We started the experiment with ∼480K cells (∼10K per well). Since cells are lost throughout the barcoding process only ∼8% of the cells were detected in the last barcoding step, before cell lysis and sub-library generation. This level of cell loss is comparable to that reported in other combinatorial barcoding experiments [25]. We generated 3 sub-libraries from a total of ∼40K ACME-dissociated cells from the two planarian species.

**Figure 3:**
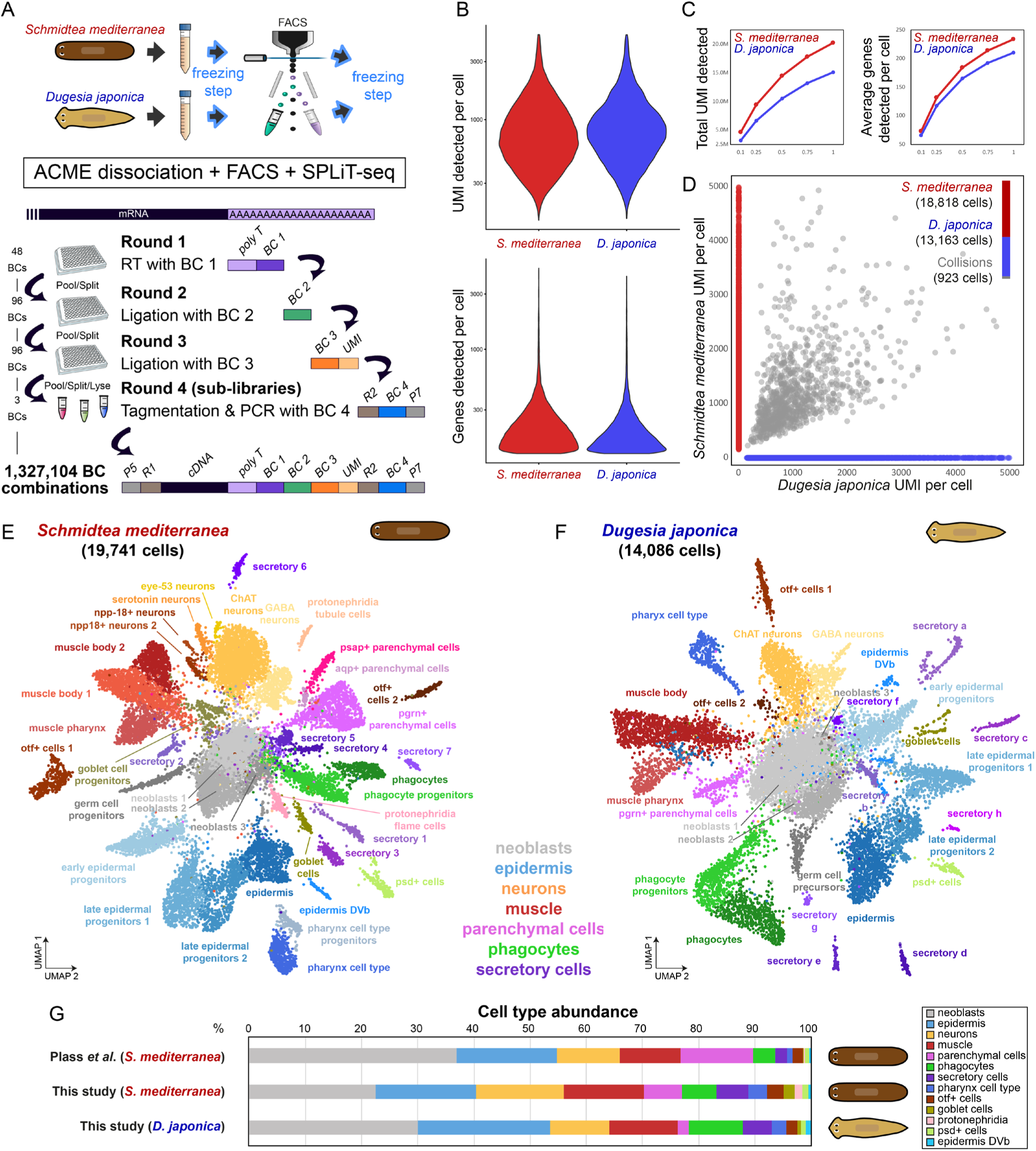
SPLiT-seq single-cell transcriptomic analysis of ACME-dissociated cells from two planarian species. **A:** Experimental workflow. We used ACME dissociated and FACS-sorted cells from the planarian species *S. mediterranea* and *D. japonica*, after two freezing steps. For SPLiT-seq, combinatorial barcoding consisted of 4 rounds of barcoding with 48 × 96 × 96 × 3 barcodes. cDNA molecules coming from each cell are uniquely labelled by one of the 1,327,104 possible barcode combinations. **B:** Violin plots showing the distribution of UMI counts and genes detected per cell. **C:** Saturation plots for UMIs per cell (left) and genes per cell (right) at given fractions of the complete sequencing depth, for the 19,741 and 14,086 single cell transcriptomes sequenced above threshold for *S. mediterranea* and *D. japonica* respectively. **D:** Scatter plot of *S. mediterranea* (red) versus *D. japonica* (blue) UMI counts per cell. Collisions shown in grey. **E-F:** UMAP visualization of 19,741 *S. mediterranea* cells (F) and 14,086 *D. japonica* cells (G), coloured by cluster identity and annotated on the basis of marker genes and homologous marker genes respectively. **G:** Comparison of cell proportions for *S. mediterranea*, in comparison with a previous cell type atlas (Plass *et al*.) and *D. japonica*. Cell clusters are grouped by cell type group.

Our cDNA sub-libraries ranged between 400-1000bp in length (Supplementary Figure 1C). We subjected these libraries to NovaSeq Illumina sequencing (paired 150 bp read length), with 425, 502 and 436 million (M) read pairs from clusters passing filter. After removal of low quality, truncated and adapter chimera sequences, read depth was 173, 192 and 196 M read pairs. To analyse this experiment we generated novel annotations (see Methods) of the most recent *S. mediterranea* genome assembly [47] and the recently sequenced *D. japonica* genome [48]. We mapped 96% and 85% of *S. mediterranea* and *D. japonica* reads to gene models in their respective gene annotations. Despite obtaining a similar amount of reads mapping to the *D. japonica* genome, the genome annotation of this species is less complete, due to the fragmented assembly of this species. We selected cell barcodes with reads mapping to ≥125 genes (19,975 and 14,263 cell barcodes in *S. mediterranea and D. japonica* respectively) and discarded excessive UMI containing cell barcodes (>5000 counts) to prevent the inclusion of multiplets (234 and 177 cell barcodes excluded in *S. mediterranea and D. japonica* respectively). This rendered 19,741 cells for *S. mediterranea* and 14,086 cells for *D. japonica*. The latter species has a comparatively less complete genome annotation, and consequently, fewer cells go through the high gene coverage threshold. Sub-library 3 contained fewer cells of both species, but those cells above thresholds in sub-library 3 were comparable to other libraries in UMI content (Supplementary Figure 1D) and had cells distributed throughout all clusters (Supplementary Figure 1E). Within these sets, after excluding high UMI cells, we obtain an average of 897.5 UMI per cell for *S. mediterranea* and 949.8 UMI for *D. japonica* (Figure 3B), a number similar to those obtained by recent combinatorial single cell transcriptomic studies by SPLiT-seq [34] or Sci-RNA-seq3 [25]. At this depth, we observe that libraries are not yet saturated, as shown by evaluating the number of UMIs and genes detected after performing subsampling at different fractions of the total depth (Figure 3C). To estimate the presence of collisions (cells sharing the same cell barcode) we took advantage of the species-mixing approach we mapped reads to a combination of both genomes. We detected only 923 (2.8%) cell barcodes with mapping to both species above a stringent cutoff of 10% (Figure 3D), showing the quality of our SPLiT-seq data.

To establish cell types and abundances in both species we further analysed our sets of 19,741 and 14,086 cells. We found 39 cell clusters in *S. mediterranea* (Figure 3E, Supplementary Table T1). Three central clusters highly expressed neoblast makers such as *Smedwi-1* (Supplementary Figure 2). Within the remaining clusters, we found cells expressing markers (Supplementary Table T2) from all progenitor and differentiated cell types that we described previously using Drop-seq [13] (Supplementary Figure 2). Some groups were clustered together in fewer clusters (Supplementary Figure 2). For instance, we identified only 3 clusters containing parenchymal cell types, also termed cathepsin+ cells [12], but within them we identify markers of all 7 clusters of parenchymal cells described in Plass *et al*. In other cases, such as the secretory cell types, we achieved better clustering (Supplementary Figure 2), finding 8 well-resolved cell clusters, compared to 4. Remarkably, we found a novel neoblast cluster containing *nanos-*positive germ cell precursor cells (Supplementary Figure 3A). These cells are well described in the literature [49, 50], but none of the previous trypsin-based planarian single cell transcriptomic studies [12, 13, 16] were able to distinguish them from the other neoblast populations (Supplementary Figure 3A). Though these germ cell precursor cells are rare (1.6% of our total cell number), single-cell methods can detect far rarer cell types. Furthermore, previous studies included more cells and/or higher UMI contents. Therefore, these facts cannot explain their clustering together with other neoblast populations in other studies. This strongly suggests that the detection of these germ cell precursors in our study relies on the early fixation provided by ACME. We detected low abundance cell clusters described in Plass *et al*., including two protonephridia cell types (tubule and flame cells, 0.4 and 1.0% respectively), psd+ cells (1.0%), epidermis of the dorsoventral boundary (DVb) (0.5%), and even lower abundance neuron subtypes including eye-53 expressing neurons (0.2%).

We then aimed to analyse for the first time the *D. japonica* cell type atlas. This planarian species belongs to another planarian clade, and its last common ancestor with *S. mediterranea* lived ∼85 million years ago [51]. Due to its comparatively lower cell numbers we were able to confidently detect fewer cell clusters. We annotated cell types by comparing their markers to their *S. mediterranea* homologues (Figure 3H, Supplementary Table T1), finding similar cell types in comparable relative abundances. While it is difficult to establish one to one homology of cell types based on top markers [52] we confidently detected the major cell type groups with this approach (Supplementary Table T2): neoblasts, epidermis, neurons, muscle, parenchymal cells, phagocytes and secretory cells. At this resolution, we are only able to cluster two types of neurons and muscle respectively. We also identify low abundance cell types, such as psd+ cells (0.8%) and epidermis DVb cells (1.0%). Encouragingly, our analysis recovers the germ cell precursors (1.5%) in this species as well (Supplementary Figure 3B).

We then compared the cell type compositions of both species. We grouped cell types in groups according to previous data [13]. We first compared the *S. mediterranea* dataset to our previously described dataset generated from trypsin dissociated cells analysed by Drop-seq (Figure 3G). Our *S. mediterranea* dataset contains ∼22% neoblasts, in line with microscopy-based estimates [42], obtained by the classic maceration technique. The most abundant cell type groups in *S. mediterranea* are epidermal, neural, muscular, and parenchymal cells, present at comparable proportions as those described. This shows that ACME dissociation robustly retrieves all cell types at comparable proportions, not introducing biases in cell type composition, and can retrieve even lowly abundant cell types. We then compared the two species. The most abundant cell types in both species are the stem cells, representing 20-35% of the total cell number. In both species the most abundant differentiation cell type groups are epidermal, neurons and muscle cells. Our *D. japonica* dataset contains considerably less parenchymal cells (1 cluster, 1.9% compared to 3 clusters with 6.8% of the total cells in *S. mediterranea*). These cell types were shown to vary with animal size [42] by microscopic observation. Future analyses, enabled by the characteristics of ACME dissociation and the multiplexing capacity of SPLiT-seq, will characterise the cell type composition of each species and the factors that underlie it. Furthermore, comparing the gene expression patterns of each cell cluster will provide insights into cell type evolution. With regards to our methodology, this proof of principle highlights the flexibility and efficiency of ACME by allowing the robust simultaneous processing of two (and potentially more) species in a single SPLiT-seq and sequencing run.

## Discussion

Here we present ACME dissociation, a new cell dissociation protocol for single cell transcriptomics. Our protocol relies on the principle of acetic acid dissociation, an approach used to dissociate cells for microscopy in past centuries but not yet applied to modern single-cell transcriptomics. The original maceration protocol was applied to relatively soft-bodied animals such as planarians and cnidarians. However, we have shown that the approach works, with slight modifications, in a broad range of animals with hard body parts such as chorions, vitelline membranes, cuticles, and shells. ACME cannot dissolve or penetrate these hard parts, but straightforward mechanical disruption is sufficient to extract cells from their acellular surroundings. We have successfully obtained dissociated cells from species belonging to all major animal groups from a wide range of habitats. It is possible that further modifications of the ACME protocol will specifically optimise the quality of cell suspensions in different organisms. We highlight the main criteria for species-specific optimisation (time of dissociation, mechanical disruption, filtering steps), but other changes might help provide the ideal dissociation conditions for each organism. Our protocol provides a robust and broadly applicable starting point for such optimisation. We also note that while ACME provides simultaneous dissociation and fixation, it is also worthwhile to consider ACME as a fixative of cells dissociated with other methods, as it preserves RNAs with high integrity, is compatible with cell staining and FACS, and provides an excellent platform for combinatorial methods of single-cell transcriptomics.

ACME is a cell dissociation approach that fixes cells while they are being dissociated. This has enormous advantages over enzymatic and mechanical approaches, as disrupting the cellular environment has effects on the cellular transcriptomic profiles that are only beginning to be realised [36-38]. Unlike nuclei extraction approaches, ACME preserves the cell cytoplasm, where most cellular mRNAs reside. Furthermore, the dissociation-fixation approach streamlines the preparation of cell suspensions for single-cell transcriptomics, with the possibility of cryopreserving cells before and/or after the FACS sorting. This will allow a range of experiments presently beyond the range of current approaches, including the collection of large sample sets, consisting of different treatments, time points and/or replicates. These can then be subject to simultaneous or sequential cryopreservation, and single-cell transcriptomics can be performed on all samples, multiplexed together in a single SPLIT-seq run. This represents a marked improvement on current workflows, helping prevent a variety of batch effects from accumulating. For instance, this could lead to clinicians preserving dissociated patient material to be later subjected to single-cell transcriptomics at a different research institution.

Similarly, enzymatic methods are hard to apply to organisms that are difficult to culture in the laboratory, as single-cell transcriptomics, or at least FACS and cryopreservation, needs to take place immediately when using previous protocols. ACME dissociation consists of simple reagents, and only requires widely available instruments: a shaker, a low speed centrifuge and a freezer. Cells are fixed from the beginning of the process and can be immediately cryopreserved. Therefore, we envision that ACME dissociation will facilitate the single-cell analysis of organisms from locations where single-cell facilities are not available, such as on field sampling trips, and allow exchange between collaborating institutions. This will accelerate our knowledge of cell types across the tree of life.

We have also shown the utility of SPLiT-seq with ACME dissociated cells. We profiled ∼34K cells of two different species in just one experiment. This large throughput in cell number is enabled by the combinatorial approach, and greatly exceeds the cell numbers obtained by nanodroplet-based approaches. UMI content per cell remains lower than those of nanodroplet-based methods, but in line with other recently published combinatorial methods. The original SPLiT-seq paper [34], using extracted nuclei, achieved an average of 1,345 UMI per nucleus in their largest library of 163K nuclei. Similarly, Cao and coworkers used Sci-RNA-seq3 and obtained an average of 671 UMI per nucleus in their large experiment of ∼2M nuclei [25]. These numbers compare well to our average UMI per cell of 897.5 for *S. mediterranea* and 949.8 for *D. japonica*. Our libraries have high complexity; therefore, higher read depth coverage would improve the UMI content of cell barcodes. Even at low sequencing depths, and with relatively low UMI counts per cell, we identify cells of all cell types that were previously described by trypsin dissociations analysed by Drop-seq [13] in the planarian *S. mediterranea* and found a similar number of cell clusters in a previously uncharacterised planarian species, *D. japonica*. This shows that the ‘many cells, few UMIs per cell’ approach is highly effective to profile cell types, as previously suggested [25], even in uncharacterised species.

Our data shows that ACME is a powerful approach with a great potential for single-cell transcriptomics by combinatorial barcoding. It is almost certain that the mRNA in ACME dissociated cells could also be analysed by other single-cell barcoding techniques such as Drop-seq or InDrop. As ACME dissociated cells are fixed and can easily be cryopreserved, we also envision that ACME could provide a framework for developing complex labelling procedures for single-cell analysis. Our current procedure involves one step of RT and two rounds of splint oligo ligation, after DNA and cytoplasm labelling. We foresee that more complex staining procedures (such as immunohistochemistry or mRNA *in situ* hybridisation, or several metabolic labelling procedures) could be used to subset cells that would be later sorted by FACS to profile lowly abundant cell populations.

Altogether, we show that ACME is a versatile and powerful cell dissociation method for single-cell transcriptomics. ACME dissociation provides a solution to various shortcomings of the single-cell transcriptomic workflow, providing early fixation of material. This same fixation will enable further development of this technique. Therefore, we believe that ACME will be a valuable tool for single-cell transcriptomics that will greatly enable the investigation of cell type diversity and dynamics in multiple different organisms presently beyond the scope of these revolutionary techniques.

## Materials and Methods

### ACME dissociation

ACME solution was prepared fresh using a 13:3:2:2 ratio of commercially sourced DNase/RNase-free distilled water, methanol, glacial acetic acid and glycerol. For each sample, between 10-30 mixed size adult planarians (cultured as previously described [13]) were added to a 15 mL Falcon tube, for a final biomass volume of ∼100-300 μL. We removed planarian water using a Pasteur pipette and added ∼100-500 μL of 7.5% N-acetyl cysteine in 1x PBS, sufficient to cover the planarians. N-acetyl cysteine helps clean planarian mucus and protects RNA (See Supplementary Note 1). The ACME solution was immediately added to samples to a final volume of 10 mL per tube. Alternatively, N-acetyl cysteine can be added to the ACME solution at this stage, in the same quantity as noted above.

Samples were left to dissociate at room temperature for 1 hour on a see-saw motion shaker at 35-45 rpm, with tubes oriented parallel to the direction of movement. We then pipetted the reactions up and down several times to complete dissociation using 1ml pipette tips. From this point, samples were kept on ice to prevent RNA degradation. We centrifuged samples at 1000 g for 5 min (4°C) to remove the ACME solution. The resulting pellet is not completely compact, so the supernatant must be discarded carefully. To clean the cells, 7 mL of buffer (1x PBS 1% BSA) was added and the pellet was mixed by flicking. Samples were centrifuged again at 1000 g for 5 min (4°C) and the supernatant was removed. If the pellet was still not compact, an additional cleaning step was performed. Pellets were resuspended in 900 μL of buffer (1x PBS 1% BSA) and transferred to 1.5 mL Eppendorf tubes.

To cryopreserve cells, we added 100 μL of DMSO per tube [43] and stored samples directly at -80°C. Afterwards we thawed samples on ice and centrifuged them at 1000 g for 5 min (4°C) to remove DMSO. The supernatant was discarded and pellets resuspended in 1 mL of washing buffer (1x PBS 1% BSA). Samples were centrifuged again and pellets again resuspended in 1 mL of fresh washing buffer.

ACME dissociation in other animals was performed with modifications to the above protocol. Zebrafish embryos were dechorionated before the ACME solution was added, and mechanically disrupted in the ACME solution by applying short pulses of Polytron homogenisation. *Lymnaea stagnalis* embryos were decapsulated by passing them through a syringe and then mechanically disrupted in the ACME solution. To remove broken eggshells, the cell dissociation mixes for both these species were passed through a 100 μm CellTrics filters (Sysmex). *Parasteatoda tepidariorum* egg capsules were either manually dissected under the scope within the ACME solution or mechanically disrupted in the ACME solution using short pulses of Polytron homogenisation, and filtered through a 40 μm cell strainer (Corning). *Pristina leidyi* adults were manually shaken every 10 minutes during dissociation.

### Flow cytometry and FACS

ACME dissociated cells were filtered through 50 μm CellTrics strainers (Sysmex), collected in 1.5 mL Eppendorf tubes and stained with the nuclear dye DRAQ5™ (eBioscience) adding 0.5 μL/mL of 5 mM stock solution, and the cytoplasmic dye Concanavalin-A conjugated with AlexaFluor 488 (Invitrogen), adding 2 μL/mL of 1 mg/mL stock solution. Staining concentrations require optimization and ultimately depend on FACS/Cytometer adjustments and cell concentration. Cells were stained in the dark, on ice, for 30-45 min, and visualized using a CytoFlex S Flow Cytometer (Beckman Coulter) or sorted using a BD FACS Aria III (BD Biosciences) Cell Sorter. To avoid RNAse contamination during cell sorting, the FACS was thoroughly decontaminated with bleach and precooled before sorting, keeping injection and collection chambers at 4°C during the process. Sorting was performed using BD FACSDiva Software, setup in 4-Way Purity mode, with an 85 μm nozzle and moderate-pressure separation (45 Psi). We set single-cells, DRAQ5 positive, Concanavalin-A positive to be sorted and collected together in 1.5 mL Eppendorf tubes with 100 μL of collection buffer (1x PBS, 1% BSA), obtaining up to 500,000 cells per tube. Completing a run normally takes 3 to 5 hours.

After sorting, samples were centrifuged at 1000 g for 5 min (4°C). The supernatant was removed and the pellet resuspended in 900 μL of fresh buffer (1x PBS 1% BSA). We cryopreserved cells at this point, by adding 100 μL of DMSO and storing them at -80°C.

### Split Pool Ligation-based Transcriptome sequencing (SPLiT-seq)

The SPLiT-seq protocol was performed as previously described [34] with modifications described below. Sorted samples were thawed, centrifuged at 1000 g for 5 min (4°C), the supernatant was removed, and the cells resuspended in 100-200 μL of buffer (1x PBS 1% BSA) and pooled together. We stained 100 uL of a 1:10 dilution of these cells for 15-20 min and counted this subsample by flow cytometry. The remaining portion of the pooled cells were then diluted to a final concentration of 1.25 M cells/mL (10,000 cells per well for the reverse transcription round).

### Plates preparation

Barcodes were provided lyophilized by Integrated DNA technologies on three 96-well Stock-Plates: Stock-1 (well-specific anchored poly(dT) Round 1 Barcodes), Stock-2 (well-specific Round 2 Barcodes) and Stock-3 (well-specific Round 3 Barcodes). Lyophilized barcodes were resuspended in DNAse/RNAse free water to a final concentration of 100 μM/well. From the Stock Plates, we prepared another three plates at specific working dilutions (WD-Plates). Using a multichannel pipette, we prepared WD-1 with 12 μL of the correspondent barcode from Stock-1 and 88 μL of DNAse/RNAse free water per well. For WD-2, we mixed 12 μL of the correspondent barcode from Stock-2, 11 μL of Linker_1 (100 μM) and 77 μL of DNAse/RNAse free water per well. Finally, WD-3 was made from 14 μL of the correspondent barcode from Stock-3, 13 uL of Linker_2 (100 μM) and 73 μL of DNAse/RNAse free water per well.

With a total volume of 100 μL per well, the WD-1 plate will last for up to 25 experiments (4 μL used per experiment), while WD-2 and WD-3 plates will last for up to 10 experiments (10 μL used per experiment). Before following the SPLiT-seq protocol, the WD-2 and WD-3 plates were heated to 95°C, for 2 min, and ramped down to 20°C at a rate of -0.1°C/s, to anneal the 5’ end of each barcode oligo to the universal linker oligos.

### Round 1 of barcoding: Reverse transcription

The first round of barcoding was carried out by in-cell reverse transcription (RT). The original SPLiT-seq protocol uses a combination of random hexamer and anchored poly(dT) oligos [34]. We only used the latter (Supplementary Table T3). A nuclease-free 96-well plate (Round-1) was prepared on ice by transferring 4 μL/well from the equivalent positions in WD-1. We then added 8 μL/well of RT mix to the plate: 4 μL of 5x RT Buffer (Thermo Scientific), 0.35 μL of SUPERase-In RNAse Inhibitor (20 U/μL, Invitrogen), 1 μL of 10 mM/each dNTPs (New England BioLab), 1.65 μL of Nuclease-free water and 1 μL of Maxima H Minus RT (Thermo Scientific). Finally, 8 μL of previously counted cells (1.25 M cells/mL) were also added to each well, giving a total volume of 20 μL/well. The Round-1 plate was incubated in a thermocycler for 30 min at 50°C, and immediately placed on ice. Individual reactions were then pooled together in a 15 mL Falcon tube, on ice, and Round-1 plate was discarded. We added 9.6 μL of 10% Triton X-100 to the pooled cells (0.1% final concentration) and centrifuged them at 1000 g for 5 min (4°C). We discarded the supernatant and resuspended the pellet in 2 mL of 1x NEB buffer 3.1 (New England Biolabs).

### Round 2 of barcoding: Ligation 1

The second round of barcoding was carried out by ligation reaction (Supplementary Table T3). A new 96-well plate (Round-2) was prepared on ice with 10 μL/well of barcodes from the equivalent positions in the previously annealed WD-2 plate. Then, 2 mL of ligation mix (500 μL of T4 Ligase Buffer 10x (NEB), 100 μL of T4 DNA Ligase (400 U/μL, NEB) and 1500 μL of Nuclease-free water) was added to the 2 mL of cells in 1x NEB buffer 3.1 and mixed thoroughly into a disposable basin. We added 40 μL/well of this ligation mix (including cells) to the Round-2 plate and covered it with an adhesive PCR plate seal. The plate was incubated in a thermocycler for 30 min at 37°C. To block Linker_1 after incubation, a blocking solution was prepared with 264 μL of Blocker_1 (26.4 μM final concentration), 250 μL of T4 Ligase Buffer 10x (NEB) and 486 μL of Nuclease-free water. After incubation, the cover of the Round-2 plate was removed and 10 μL of blocking solution were added to each well, giving a final volume of 60 μL/well. The plate was sealed again, with a new adhesive cover, and incubated for another 30 min at 37°C to bind Linker_1 and Blocker_1 oligos.

### Round 3 of barcoding: Ligation 2

The third round of barcoding was carried out by a second ligation reaction (Supplementary Table T3). To prepare the Round-3 plate, we filled a 96-well plate with 10 μL/well of barcodes from equivalent positions in the pre-annealed WD-3 plate. After taking the Round-2 plate from the incubator, the cover was discarded and cells were pooled together into a new disposable basin. We added 100 μL of T4 DNA Ligase (400 U/μL, NEB) to the basin and mixed thoroughly with the cells. The Round-3 plate was then filled with 50 μL/well of this cell-ligase solution, sealed with an adhesive PCR plate seal and incubated for 30 min at 37°C. Termination solution was prepared with 288 μL of Blocker_2 (11.5 μM final concentration), 625 μL of EDTA (to stop ligase activity) and 1587 μL of nuclease-free water. We added 20 μL/well of the termination solution to the Round-3 plate (for a final volume of 70 μl/well) without further incubation. Afterwards, cells were pooled into a 15 mL Falcon tube and placed on ice.

### Cell lysis

Following the addition of 70 μL of 10% Triton-X 100 (0.1% final concentration), the pooled cells were centrifuged at 1000g for 5 min (4°C). We carefully removed the supernatant (leaving about 100 μL) and resuspended the cells in 4.04 mL of washing buffer (4 mL of 1x PBS and 40 μL of 10% Triton X-100, premixed to allow Triton bubbles to settle). The cells were centrifuged again at 1000 g for 5 min (4°C). After removing the supernatant, we resuspended the cells in 50 μL of 1x PBS buffer. We diluted 5 μL of this resuspension in 195 μL of 1x PBS and counted the number of cells by flow cytometry to decide the number of sub-libraries. Then, we aliquoted the remaining 45 μL of cells in 1.5 mL Eppendorf tubes according to the cell concentration obtained by flow cytometry. For the experiment described here, we generated three different sub-libraries of ∼15.000 cells/each. The volume of each sub-library was adjusted to 50 μL with 1x PBS.

Lysis buffer was prepared with Tris pH 8.0 (20 mM), NaCl (400 mM), EDTA pH 8.0 (100 mM) and SDS (4.4%). We added 50 μL of lysis buffer and 10 μL of Proteinase K (20 mg/mL) to each sub-library and incubated the lysates at 55°C for 2 hours, shaking the tube manually every 15 min. After incubations, lysates were frozen at -80°C.

### cDNA purification with magnetic beads

We used 44 μL of Dynabeads™ MyOne™ Streptavidin C1 (Invitrogen) per lysate to purify the cDNA by linking the beads to the biotin molecule at the 3’ end of the third barcode. We followed the manufacturer’s protocol for Dynabead nucleic acid purification, with modifications taken from Rosenberg et al’s protocol: Manufacturer’s Washing Buffer (1x B&W) was prepared with the addition of 0.05% final concentration of Tween-20. Lysates were incubated for 10 min at room temperature with 5 μL of 100 uM PMSF (diluted in isopropanol) to inhibit Proteinase K activity. Lysates were incubated with the magnetic beads for 60 min, at room temperature, with rotation. During washing steps, samples were agitated in 1x B&W/Tween-20 buffer for 5 min at room temperature. Finally, beads were resuspended in 250 μL of 10 mM Tris-T buffer (10 mM Tris-HCl pH, 0.1% Tween-20 and 0.2% SUPERase-In RNAse Inhibitor) and kept at 4°C.

### Template Switch

The template switch mix was prepared with 44 μL of 5x RT Buffer (Thermo Scientific), 44 μL of 20% Ficoll PM 400 (Sigma Aldrich), 22 μL of 10 mM/each of four dNTPs (NEB), 5.5 μL of TSO primer (100 μM), 11 μL of Maxima H Minus RT and 93.5 μL of nuclease-free water per sample. Beads linked to cDNA were washed, using a magnetic rack, with 250 μL of nuclease-free water (no resuspension) and resuspended in 200 μL of the template switch mix. Samples were incubated in the mix for 30 min at room temperature and then for 90 min at 42°C, with agitation. After incubation, the template switch mix was removed using a magnetic rack; beads were resuspended in 250 μL of Tris-T buffer and kept at 4°C. At this point, beads can be kept in the fridge (4°C) for up to 4 days and be reused at least once for subsequent steps.

### PCR amplification

The PCR mix was prepared with 110 μL of 2x Kapa HiFi HotStart ReadyMix (Roche), 8.8 μL of PCR_PF (10 μM), 8.8 μL of PCR_PR (10 μM) and 92.4 μL of nuclease-free water. Samples from the template switch reaction were placed in a magnetic rack and washed with 250 μL of water (no resuspension). After removal of wash fluid, they were then resuspended in 220 μL of the PCR mix and each split into 4 PCR tubes. The following program was run in the thermocycler: 95°C (3 min) and five cycles at 98°C (20 s), 65°C (45 s) and 72°C (3 min). The 4 reactions for each library were then combined in single 1.5 mL Eppendorf tubes per library, and Dynabeads were separated using a magnetic rack. 200 μL of supernatant, containing cDNA in suspension, were split into 4 PCR tubes once again, and transferred to a qPCR plate (50 μL/well). We added 2.5 μL of 20x EvaGreen (Biotium) to each well and run the following program in a qPCR thermocycler: 95°C (3 min), cycling until plateau phase, normally 8-10 cycles, at 98°C (20 s), 65°C (20 s) and 72°C (3 min), and a final elongation at 72°C (5 min).

### Size selection

We purified qPCR reactions by SPRI size selection to remove fragments smaller than 300 bp. We used Kapa Pure Beads (Roche) at a ratio of 0.8x and followed the manufacturer’s protocol for “Cleanup of Fragmented DNA in NGS Workflows”, with two modifications taken from the original SPLiT-seq protocol: washing steps were performed with 750 μL of 85% ethanol and cDNA was eluted in 20 uL of nuclease-free water at 37°C for 10 min.

### Tagmentation

The sub-libraries were tagmented using the Nextera DNA Library Preparation Kit (Illumina). After SPRI 0.8x size selection described above, we quantified the sub-libraries by Qubit (Thermo Fisher) and diluted 50 ng of cDNA in a total volume of 20 μL of nuclease-free water. The tagmentation reaction mix was prepared with 20 μL of cDNA (50 ng), 25 μL of Tagmentation Buffer and 5 μL of Enzyme 1, and incubated in a pre-heated thermocycler for 5 min at 55°C, placed immediately on ice at the end of this time period. We neutralized the tagmentation activity of Enzyme 1 by immediately cleaning the reaction with the Monarch PCR & DNA Cleanup Kit (NEB) according to the manufacturer’s protocol. Samples were eluted in a final volume of 20 μL of UltraPure water.

### Round 4 of barcoding: PCR

The fourth barcode was introduced by PCR (Supplementary Table T3). We prepared a separate reaction mix for each library, containing 20 μL of tagmented cDNA, 25 μL of 2x Kapa HiFi HotStart ReadyMix (Roche), 1.5 μL of P5_oligo (10 μM) and 1.5 μL of Round 4 Barcode (10 μM). For the three sub-libraries used in the present study, we used the Round4_1, Round4_2 and Round4_3 oligos. Parallel to the PCR reaction, we ran a qPCR by adding 2.5 μL of 20x EvaGreen (Biotium) to one replica sample to visualize the plateau phase. The PCR program ran as follows: 95°C (30 s), cycling until plateau phase (8-10 cycles) at 95°C (10 s), 55°C (30 s) and 72°C (30 s), and final elongation at 72°C (5 min). PCR reactions were size selected (SPRI 0.7x) by mixing 40 μL of sample with 28 μL of Kappa Pure Beads (Roche), and following the size selection protocol described above. The sub-libraries were each resuspended to a final volume of 20 μL in nuclease-free water and fragment distribution was checked running a bioanalyzer (Agilent High Sensitivity DNA Kit) according to the manufacturer’s protocol.

### Sequencing and Quality Control

The three sub-libraries were pooled together and sequenced on a NovaSeq 6000 platform (Illumina) by Novogene, with 150bp length, paired end reads. These reads were provided without any quality verification except a basic chastity check. They were therefore subject to initial quality checks with FastQC. Read Phred quality was generally good, but adaptor and N content required curation and removal. CutAdapt v2.8 [53] was used to trim residual adaptor sequence, low quality, and short reads. Differing strategies for clean-up were used for Read 1 (transcript sequence) and Read 2 (UMI and barcode sequences). For read 1, cutadapt -j 4 -m 60 -q 10 -b AGATCGGAAGAG was run, removing residual Illumina universal adapter and a read length shorter than 60 bp. For read 2, cutadapt -j 4 -m 94 --trim-n -q 10 -b CTGTCTCTTATA was run, removing reads shorter than 94 bp (the minimum to span all barcodes), terminal Ns, and residual Nextera adapter sequence. Read 2 sequences were checked for “phase” (i.e., whether barcodes were in their correct position, due to possible indels) using grep to compare adapter-derived flanking sequence was correctly positioned with that of each read. Reads were conservatively retained, with only reads with UMI and UBC barcodes in the correct location carried forward to further analysis. Dephasing, while advisable, did not prove a major issue, and very few reads were discarded. Makepairs (https://github.com/sestaton/Pairfq/wiki/makepairs) was used to retain only paired reads, and a further round of FastQC analysis was used to confirm all detectable adaptor and low quality sequence had been removed.

### Read mapping, barcode extraction and matrix production

The *S. mediterranea* S2F2 genome [47] was downloaded from Planmine, and the *D. japonica* v 1.0 genome [48] was downloaded from http://www.planarian.jp. *De novo* gene models were created for both *S. mediterranea* and *D. japonica*. 183 published *S. mediterranea* and 43 *D. japonica* RNAseq datasets were downloaded from the NCBI SRA and the DNA Data Bank of Japan, comprising all those listed at the time of analysis. These collected reads were aligned to the respective reference genomes using HiSat 2.1.0 [54]. StringTie and StringTie—merge [55] were then used to merge mapping outputs with the existing SMESG-high confidence gene models from Planmine (*S. mediterranea*) and the full v1 AUGUSTUS-derived gene models from http://www.planarian.jp (*D. japonica*). Isoformal variants whose length was greater than 100 kb were removed from the gene set as likely artefacts (588 in *D. japonica*, 617 in *S. mediterranea). D. japonica* and *S. mediterranea* fasta and gtf files were then concatenated to create a combined database for mapping. Drop-seq_tools-2.3.0 (https://github.com/broadinstitute/Drop-seq) was then used to create sequence dictionary, refFlat, reduced GTF and interval files. A STAR-2.7.3a [56] index was generated using the --sjdbOverhang 99 --genomeSAindexNbases 13 --genomeChrBinNbits 14 settings for both genomes concatenated together (to allow measurement of collisions, see below).

Each of the three sub-libraries sequenced in the present manuscript was then processed separately. SPLiTseq toolbox (https://github.com/RebekkaWegmann/splitseq_toolbox), which incorporates many of the components of Drop-seq_tools-2.3.0 (https://github.com/broadinstitute/Drop-seq) was used to extract, check and correct barcodes (corrections with hamming distance ≤ 1). Mapping was performed using STAR-2.7.3a [56], with --quantMode GeneCounts and all other default settings. Picard v2.21.1-SNAPSHOT (Broad Institute, http://broadinstitute.github.io/picard/) SortSam and MergeBamAlignment were used to re-order and merge aligned and tagged reads. Drop-seq_tools-2.3.0 TagReadWithInterval and TagReadWithGeneFunction were then run sequentially to note mapping location, using the custom refFlat and genes.intervals files created above. An additional character (A, T or C) was then added to the cellular barcode of each sub-library, to allow identical cell barcodes from different sub-libraries to be differentiated. Reads mapping to *D. japonica* and *S. mediterranea* were then separated. These mapping files were then used to create expression matrices using Drop-seq_tools-2.3.0 DigitalExpression for each library individually, with the following settings: READ_MQ=0, EDIT_DISTANCE=1, MIN_NUM_GENES_PER_CELL=100, LOCUS_FUNCTION_LIST=INTRONIC. These matrices, along with the novel gene models and raw reads, have been uploaded to the NCBI GEO, accession GSE150259.

### Seurat feature identification and clustering

The digital expression matrices for each sub-library were loaded into Seurat v 3.1.0 [57] within R v 3.6.2 and a Seurat object created for each, with min.cells = 1 and min.features = 125 (although note the matrix creation cutoff described above). Cells with a UMI count greater than 5000 were excluded with subset = nCount_RNA < 5000. Data was normalised, and variable features selected (selection.method = “vst”, nfeatures = 10000) and data scaled. The three libraries were then merged (merge (x = L1, y = list(L2, L3), add.cell.ids = add.cell.ids, merge.data = FALSE)). Data was again normalised (normalization.method = “LogNormalize”, scale.factor = 10000) and variable features selected (split.by = “library”, nfeatures = 10000, verbose = TRUE, fvf.nfeatures = 10000, selection.method = “vst”). Principal component analysis (PCA) was run for 50 principal components. JackStrawPlots and ElbowPlots were generated to understand the dimensionality of the data (plots not shown). Clustering was performed with resolution = 1 (*D. japonica)* or 1.4 (*S. mediterranea)* and the original Louvain algorithm. UMAP representations were created with the following settings: dims = 1:50, reduction = “pca”, spread = 1, metric = “euclidean”, seed.use = 1, and n.neighbors = 45, min.dist = 0.4 (*D. japonica)* and n.neighbors = 35, min.dist = 0.5 (*S. mediterranea)*. UMAP projections were displayed both by cluster and by library to assay for batch effects. Further batch effect correction was not necessary. A number of resolutions were trialled and compared manually with marker lists to determine the most appropriate cutoff for display. Clusters were recoloured natively within Seurat as plots were generated. Markers were extracted from Seurat using the FindAllMarkers function. Cell numbers per cluster were extracted using the Idents function. FeaturePlot was used to highlight expression of individual markers in cells in our dataset, to aid with cell cluster identification. VlnPlot was used to generate violin plots of gene and UMI content in each library, using a log scale.

### Cluster identity assignment

To establish the identity of *S. mediterranea* cell clusters we cross-referenced the markers found in them with the markers from our previous publication [13]. Representative examples are shown in Supplementary Figure S2. Novel cell clusters eye-53 neurons, serotonin neurons, protonephridia tubule cells and protonephridia flame cells were named respectively by the expression of markers *eye-53-1* (SmMSTRG.4014, dd_Smed_v6_889_0_1) [58], *sert* (SmMSTRG.6717, dd_Smed_v6_12700_0_1) [59, 60], *CAVII-like* (SmMSTRG.5392, dd_Smed_v6_4841_0_1) [61] and *egfr-5* (SmMSTRG.13890, dd_Smed_v6_11310_0_1) [62]. To establish the identity of *D. japonica* cell clusters we found known *S. mediterranea* homologues of the top *D. japonica* cluster markers as noted below, and examined the expression of these in our *S. mediterranea* and *D. japonica* feature maps.

To establish marker homology between our novel gene models and known gene sequences from previous publications we used a different approach for each species. For *S. mediterranea* comparisons to previously catalogued genes: blastn megablast [63] of known nucleotide sequences to our gene models (*E* value cutoff = *E* < 10^−99^, although best hit normally = 0) and Standalone BLAT v. 36×5 (-out=blast8) [64] were used to find clearly homologous sequences. For *D. japonica* the same approach was used to find homologues between our novel gene models and known *D. japonica* genes when appropriate. However, to annotate based on known *S. mediterranea* sequences, the protein sequences of previously identified *S. mediterranea* genes were used to search our novel *D. japonica* gene set using tblastn (*E* value cutoff = *E* < 10^−99^), alongside nucleotide vs nucleotide blat searches as described previously. Secretory clusters were arbitrarily named 1-7 and a-h according to their abundance, as the exact correspondence of these with previously published experiments and the homology between planarian species requires further research.

### Other analyses performed for single cell RNAseq data

To assay whether our library sequencing was saturated at the given read depth, reads were downsampled randomly using seqtk (10%, 25%, 50% and 75% of total read depth). These read results were extracted from our final bam files and used to generate UMI and gene numbers per cell for the cell barcode set identified from our full results (NB: as bam files were ordered in the process of data generation, sampling from the bam file directly would likely have returned non-random results). Using the matrices produced from the full data for *S. mediterranea* and *D. japonica*, a 2 species “barnyard” plot was built, showing the UMIs per cell barcode from each of the species. Collisions were defined as cell barcodes sharing over 10% of their UMIs with the minority species in this plot. Seurat’s FeaturePlot function was used to colour cells within our plots expressing particular gene markers.

## Supplementary Note 1: Protecting RNA from degradation

In the course of ACME dissociation RNAs are exposed to hydrolysis or degradation by RNAses. Therefore, RNA protective, RNAse-free conditions are essential. We have found that N-Acetyl-L-Cysteine (NAC) results in better RNA integrity after maceration. NAC was initially added as a mucolytic agent to planarian ACME dissociations. NAC is a reducing agent that breaks up disulphide bonds in the mucus, therefore solubilising it. NAC is also widely used due to its antioxidant properties [44]. NAC is acidic in solution and can be therefore mixed with the acidic ACME solution, resulting in reducing, RNA protective, conditions. We first used NAC as an initial wash step. Due to its acidity, this step has to be performed quickly as it results in cell dissociation and lysis. We also have had good results mixing the NAC directly with ACME solution. The concentration of NAC in the ACME solution and the possibility of performing it as an initial wash step should be evaluated in each case.

We have found that a significant source of RNAse contamination can come from the use of BSA. This is used to avoid cell clumps. While RNAse-free BSA can be obtained commercially, it is often too expensive to be used in large amounts. We have had good results with commercial non RNAse-free BSA. It is essential to assay its RNAse activity in advance, as this can vary considerably. We also aliquot it to prevent RNAse cross-contamination or bacterial growth. We regularly use a concentration of 1% in our protocols, but the concentration of BSA can be reduced if significant RNAse activity is present.

### Supplementary Figures

**Supplementary Figure S1:**
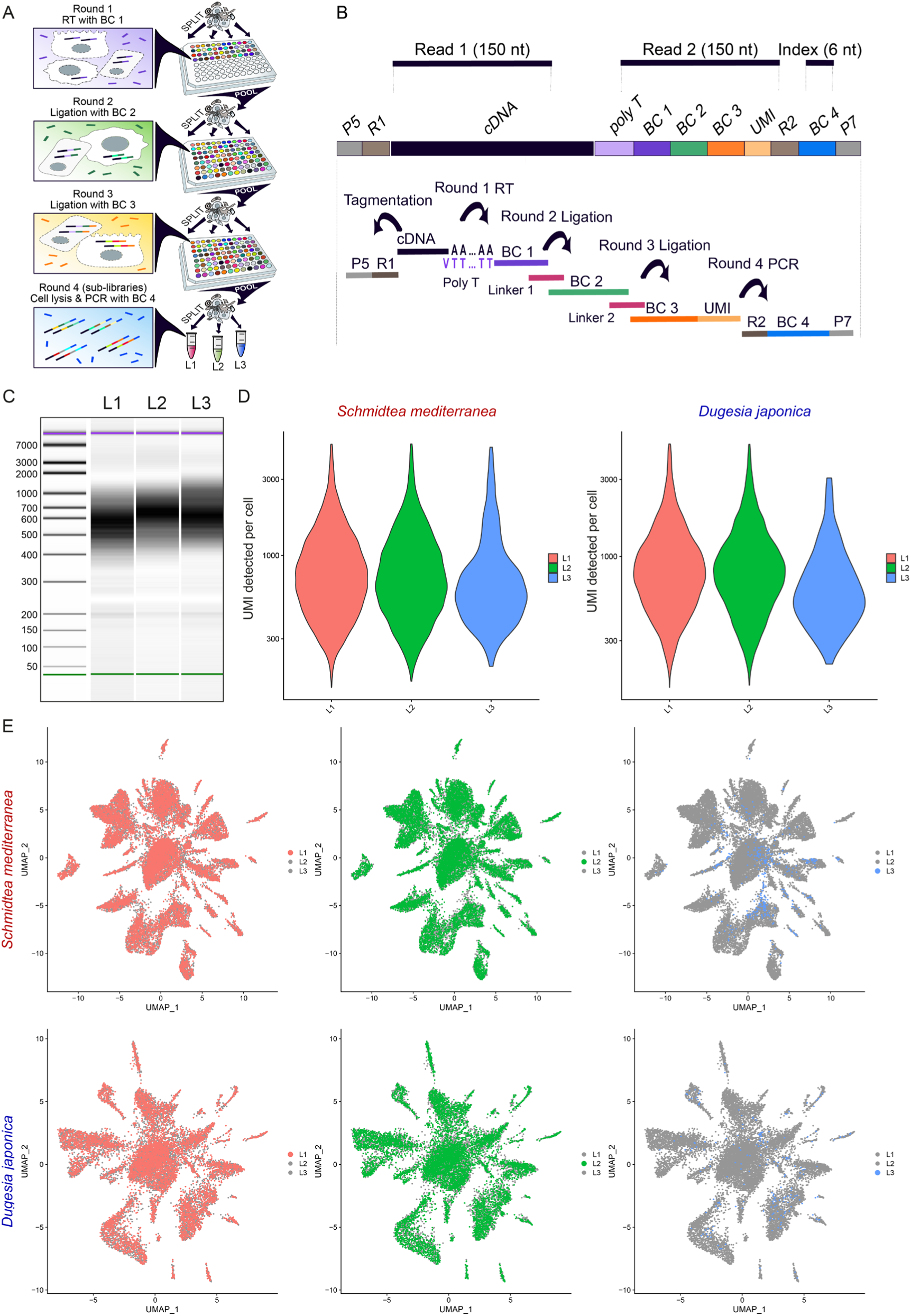
SPLiT-seq sub-library description. **A:** SPLiT-seq workflow: cells are randomly split in 96-well plates containing unique barcodes. The first RT round captures mRNAs using an anchored-poly dT barcode (BC 1). The second (BC 2) and third barcodes (BC 3, which includes the UMI) are ligated to the cDNA in two subsequent reactions. The last barcode (BC 4) is added in a final PCR reaction after tagmentation. **B:** Structure of the SPLiT-seq library. Paired-end reads (Read 1 and Read 2) and index were sequenced as indicated. **C:** Bioanalyzer profile of the 3 individual sub-libraries described here. **D:** Violin plots showing the distribution of UMI counts (left) and features detected (right) per cell in each sub-library. **E:** UMAP visualization of *S. mediterranea* cells (top) and *D. japonica* cells (bottom), coloured by sub-library.

**Supplementary Figure S2:**
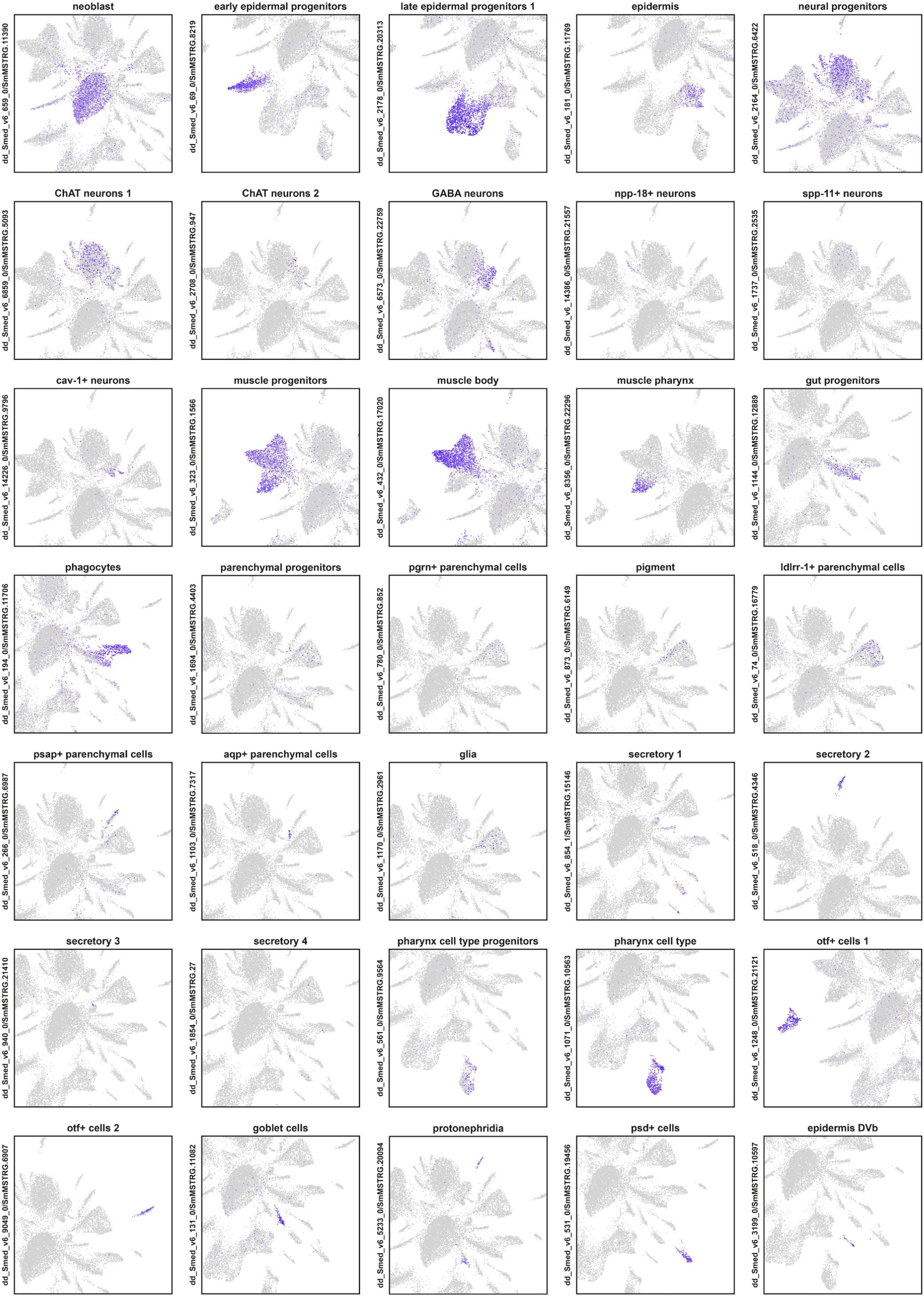
Expression of previously published cell type markers in our dataset. Feature plot of cell type markers from Plass *et al* in the UMAP visualization of *S. mediterranea* cells

**Supplementary Figure S3:**
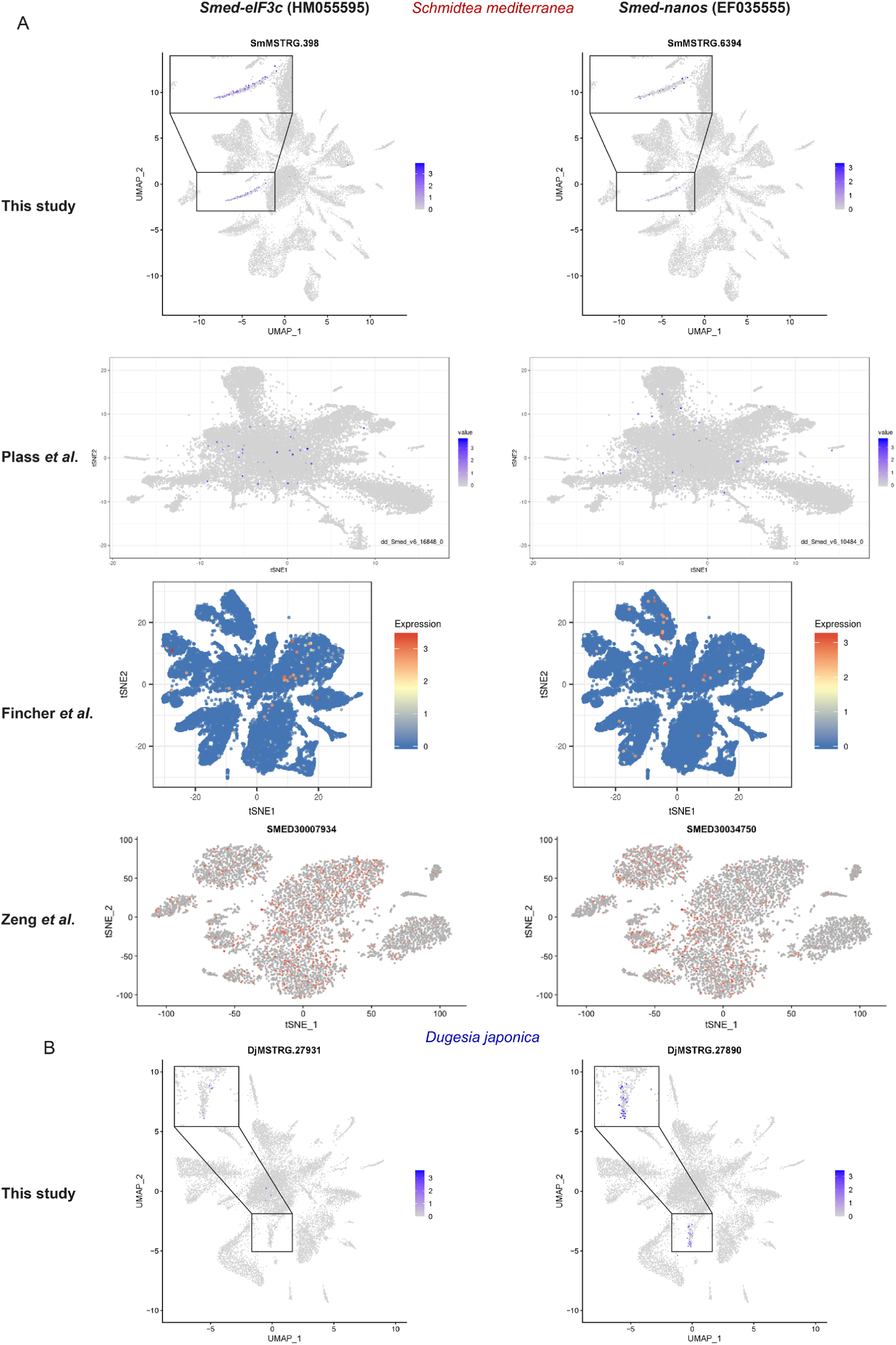
Expression of germ line precursor markers. **A:** Feature plots of germ line precursor markers *Smed-eIF3c* and *Smed-nanos* in our *S. mediterranea* dataset and previously published studies Plass *et al*. (downloaded from https://shiny.mdc-berlin.de/psca/), Fincher *et al*. (downloaded from https://digiworm.wi.mit.edu/) and Zeng *et al*. (downloaded from https://planosphere.stowers.org). **B:** Feature plots of germ line precursor markers in the *D. japonica* dataset

### Supplementary Tables

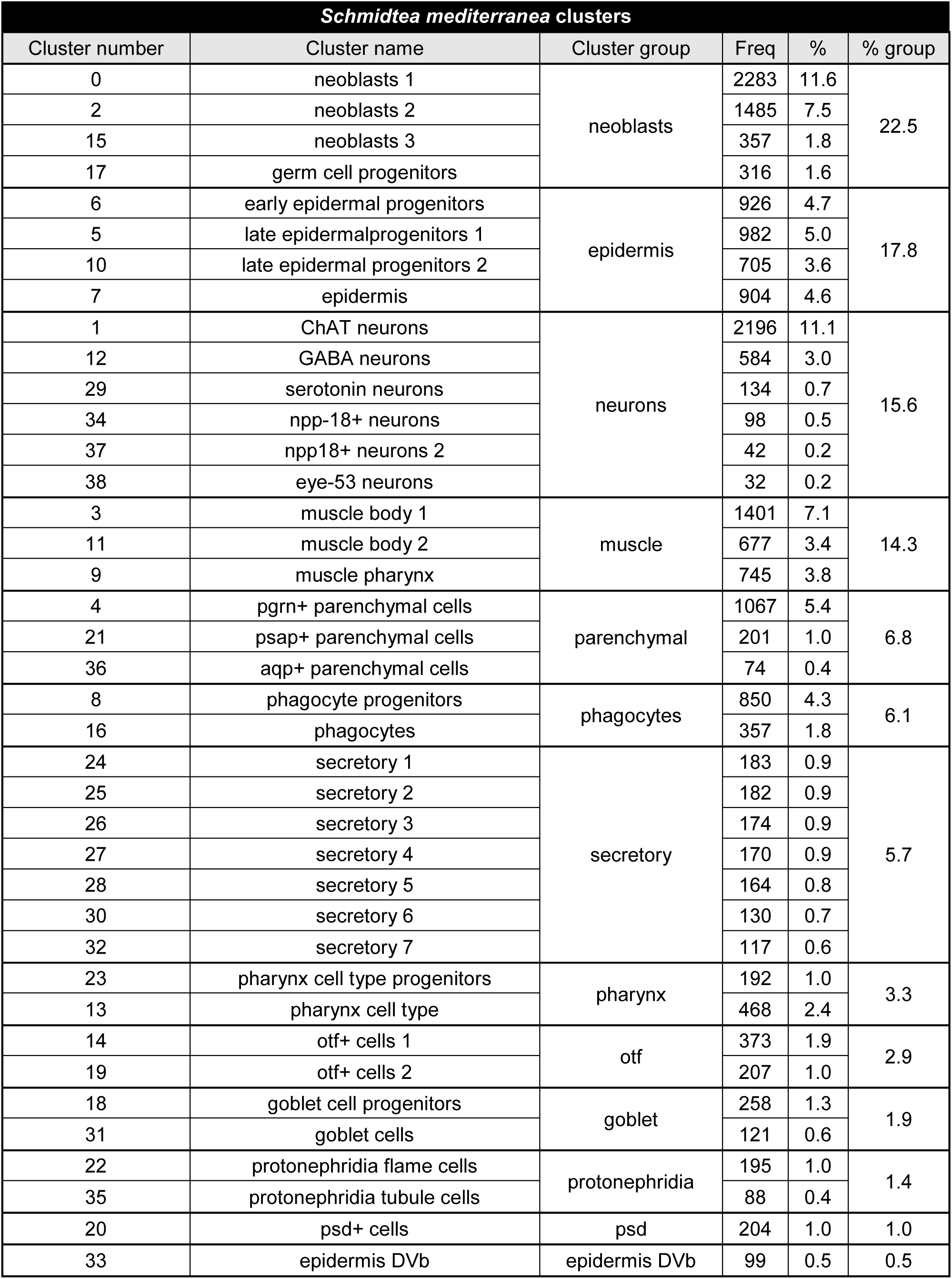
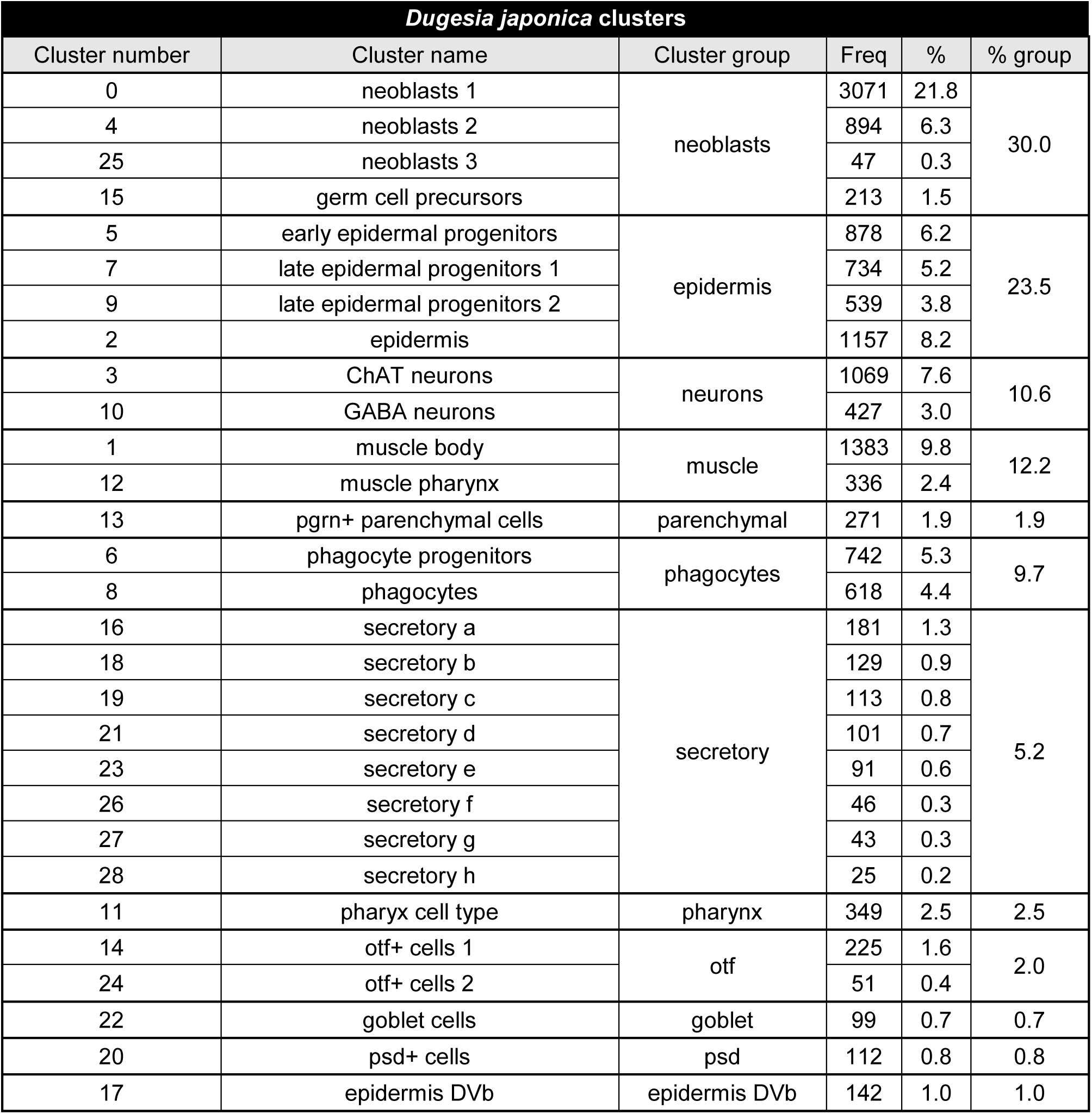

| *Schmidtea mediterranea* clusters |  |  |  |  |  |
| --- | --- | --- | --- | --- | --- |
| Cluster number | Cluster name | Cluster group | Freq | % | % group |
| 0 | neoblasts 1 | neoblasts | 2283 | 11.6 | 22.5 |
| 2 | neoblasts 2 |  | 1485 | 7.5 |  |
| 15 | neoblasts 3 |  | 357 | 1.8 |  |
| 17 | germ cell progenitors |  | 316 | 1.6 |  |
| 6 | early epidermal progenitors | epidermis | 926 | 4.7 | 17.8 |
| 5 | late epidermalprogenitors 1 |  | 982 | 5.0 |  |
| 10 | late epidermal progenitors 2 |  | 705 | 3.6 |  |
| 7 | epidermis |  | 904 | 4.6 |  |
| 1 | ChAT neurons | neurons | 2196 | 11.1 | 15.6 |
| 12 | GABA neurons |  | 584 | 3.0 |  |
| 29 | serotonin neurons |  | 134 | 0.7 |  |
| 34 | npp-18+ neurons |  | 98 | 0.5 |  |
| 37 | npp18+ neurons 2 |  | 42 | 0.2 |  |
| 38 | eye-53 neurons |  | 32 | 0.2 |  |
| 3 | muscle body 1 | muscle | 1401 | 7.1 | 14.3 |
| 11 | muscle body 2 |  | 677 | 3.4 |  |
| 9 | muscle pharynx |  | 745 | 3.8 |  |
| 4 | pgrn+ parenchymal cells | parenchymal | 1067 | 5.4 | 6.8 |
| 21 | psap+ parenchymal cells |  | 201 | 1.0 |  |
| 36 | aqp+ parenchymal cells |  | 74 | 0.4 |  |
| 8 | phagocyte progenitors | phagocytes | 850 | 4.3 | 6.1 |
| 16 | phagocytes |  | 357 | 1.8 |  |
| 24 | secretory 1 | secretory | 183 | 0.9 | 5.7 |
| 25 | secretory 2 |  | 182 | 0.9 |  |
| 26 | secretory 3 |  | 174 | 0.9 |  |
| 27 | secretory 4 |  | 170 | 0.9 |  |
| 28 | secretory 5 |  | 164 | 0.8 |  |
| 30 | secretory 6 |  | 130 | 0.7 |  |
| 32 | secretory 7 |  | 117 | 0.6 |  |
| 23 | pharynx cell type progenitors | pharynx | 192 | 1.0 | 3.3 |
| 13 | pharynx cell type |  | 468 | 2.4 |  |
| 14 | otf+ cells 1 | otf | 373 | 1.9 | 2.9 |
| 19 | otf+ cells 2 |  | 207 | 1.0 |  |
| 18 | goblet cell progenitors | goblet | 258 | 1.3 | 1.9 |
| 31 | goblet cells |  | 121 | 0.6 |  |
| 22 | protonephridia flame cells | protonephridia | 195 | 1.0 | 1.4 |
| 35 | protonephridia tubule cells |  | 88 | 0.4 |  |
| 20 | psd+ cells | psd | 204 | 1.0 | 1.0 |
| 33 | epidermis DVb | epidermis DVb | 99 | 0.5 | 0.5 |

Supplementary Table T1: Cell numbers by cluster and cell type groups
| *Dugesia japonica* clusters |  |  |  |  |  |
| --- | --- | --- | --- | --- | --- |
| Cluster number | Cluster name | Cluster group | Freq | % | % group |
| 0 | neoblasts 1 | neoblasts | 3071 | 21.8 | 30.0 |
| 4 | neoblasts 2 |  | 894 | 6.3 |  |
| 25 | neoblasts 3 |  | 47 | 0.3 |  |
| 15 | germ cell precursors |  | 213 | 1.5 |  |
| 5 | early epidermal progenitors | epidermis | 878 | 6.2 | 23.5 |
| 7 | late epidermal progenitors 1 |  | 734 | 5.2 |  |
| 9 | late epidermal progenitors 2 |  | 539 | 3.8 |  |
| 2 | epidermis |  | 1157 | 8.2 |  |
| 3 | ChAT neurons | neurons | 1069 | 7.6 | 10.6 |
| 10 | GABA neurons |  | 427 | 3.0 |  |
| 1 | muscle body | muscle | 1383 | 9.8 | 12.2 |
| 12 | muscle pharynx |  | 336 | 2.4 |  |
| 13 | pgrn+ parenchymal cells | parenchymal | 271 | 1.9 | 1.9 |
| 6 | phagocyte progenitors | phagocytes | 742 | 5.3 | 9.7 |
| 8 | phagocytes |  | 618 | 4.4 |  |
| 16 | secretory a | secretory | 181 | 1.3 | 5.2 |
| 18 | secretory b |  | 129 | 0.9 |  |
| 19 | secretory c |  | 113 | 0.8 |  |
| 21 | secretory d |  | 101 | 0.7 |  |
| 23 | secretory e |  | 91 | 0.6 |  |
| 26 | secretory f |  | 46 | 0.3 |  |
| 27 | secretory g |  | 43 | 0.3 |  |
| 28 | secretory h |  | 25 | 0.2 |  |
| 11 | pharynx cell type | pharynx | 349 | 2.5 | 2.5 |
| 14 | otf+ cells 1 | otf | 225 | 1.6 | 2.0 |
| 24 | otf+ cells 2 |  | 51 | 0.4 |  |
| 22 | goblet cells | goblet | 99 | 0.7 | 0.7 |
| 20 | psd+ cells | psd | 112 | 0.8 | 0.8 |
| 17 | epidermis DVb | epidermis DVb | 142 | 1.0 | 1.0 |

**Supplementary Table T1: Cell numbers by cluster and cell type groups**

**Supplementary Table T2: Top markers by cluster**

(see supplementary excel file)

**Supplementary Table T3: List of Barcodes**

(see supplementary excel file)

## Data Availability

The data that support the findings of this study are available from GEO Accession GSE150259. This includes all sequencing reads (SRA Accession SRP261093), reannotated gene models for both species analysed, and the final gene expression matrices used for analysis. These data, as well as further resources, are also available from https://jakke-neiro.github.io/Oxplatys/

## Acknowledgments

This work was supported by an MRC grant (MR/S007849/1) and a Royal Society Grant (RGS\R1\191278) to JS. HG-C was supported by a Nigel Groome studentship from Oxford Brookes University. PA-C was supported by an EMBO Long Term Fellowship (ALTF-217-2018). JN was supported by funding from a BBSRC grant (BB/M011224/1) and the Osk. Huttunen Foundation (Doctoral grant). We thank Helen Ferry and Liam Hardy at the Experimental Medicine Division Flow Cytometry Facility at the Nuffield Department of Clinical Medicine (University of Oxford), Michal Maj at the Flow Cytometry Facility at the Dunn School of Pathology (University of Oxford) and Òscar Fornas at the Centre for Genomic Regulation/Universitat Pompeu Fabra FACS Unit (Barcelona) for their help and advice with Flow Cytometry. We thank Alistair McGregor at Oxford Brookes University for providing useful comments and discussion about the dissociation, Teresa Adell of the Department of Genetics, University of Barcelona and Christopher Laumer at the European Bioinformatics Institute (Cambridge) for discussion and advice on the protocol. We thank Mireya Plass at the Centre for Regenerative Medicine of Barcelona for computational analysis advice and for critical review of the manuscript.

